# Shining a light on daytime coral spawning synchrony across oceans

**DOI:** 10.1101/2025.03.23.643993

**Authors:** Charlotte Moritz, Serge Andrefouet, Claire-Sophie Azam, Cecile Berthe, Manon Fourriere, Adeline Goyaud, Melina Grouazel, Gilles Siu, Marguerite Taiarui, Anne-Marie Trinh, Vetea Liao

**Author notes:** **Classification** Research article. **Competing interests** The authors declare no competing interests.

## Abstract

**Aim:** The variety of coral taxa and environmental cues triggering broadcast spawning gave rise to contrasting theories about coral reproduction synchrony. Here, we shine a new light on this synchrony across oceans by highlighting how environmental cues modulate spawning time at various spatial scales in an understudied yet abundant gonochoric coral species.

**Location.** South Indian and Pacific Oceans.

**Time period.** 2014-2023.

**Major taxa studied.** *Porites rus*.

**Methods:** *Porites rus* daytime spawning was investigated using a non-invasive citizen science approach (>300 voluntary observers) at colony, reef, island and ocean scales. Spawning time was recorded between 2014 and 2023 at a total of 104 reef locations from 15 islands in 3 countries and multiple depths across the Indian and Pacific Oceans. Statistical models were employed to assess the relationships between spawning time and depth, light, and sea surface temperature at different spatial scales, and in both male and female colonies.

**Results:** Spawning occurred synchronously for colonies located a few meters to >15,000 km apart, monthly five days after full moon over an extended, uninterrupted period from October to April. Strong linear relationships between depth, light, water temperature and spawning time after sunrise hold at the different spatial scales, for both males and females which spawn *ca.* 20 minutes apart. Interestingly, single colonies spawned across consecutive days and months.

**Main conclusions:** The largest dataset for a daytime coral species compiled here allows extremely accurate predictions of *P. rus* spawning months, days and time (to the minute) at different locations and depths in the Southern Hemisphere. Previously underexplored, the highly effective reproductive strategy of *P. rus* may explain its broad distribution and persistence in stressed environments, positioning it as an invaluable model organism for studying the physiological and genetic processes driving behavioural synchrony and biological rhythms across interconnected biogeographical regions.

## Introduction

Accurate synchronisation of conspecific behaviour benefits the maintenance of metapopulations through the completion of mutual goals, especially reproduction (Ims, 1990). Across interconnected marine regions, sessile corals use synchronous mass coral spawning to produce abundant offspring to be dispersed across reefs, maintaining sustainable metapopulation levels and genetic diversity worldwide. The biological clock for reproduction of most corals is set at specific seasons and days of the year (Baird, Guest & Willis, 2009) when male, female or hermaphrodite broadcast spawning corals produce gametes that use seawater to gather and mix, favouring fertilisation of female eggs by sperm cells. This simultaneous coral gamete release on a given reef or series of reefs creates a floating marine snow (bundles or female eggs) or dense smoky clouds (sperm) that fog the sea for several minutes to hours and can even be observed from space (Yamano, Sakuma & Harii, 2020). While the production and release of these cells must synchronise among nearby colonies of the same species to ensure successful genetic mixing (Kitanobo, Toshino & Morita 2022), a species-specific timing (i.e. different coral species reproduce at various times of the year, lunar cycles, and day or night times) also avoids coral species mixing and favours speciation (Fukami et al. 2003, Levitan et al. 2011, Baird et al. 2022).

Several environmental cues were identified to influence synchronous coral gamete maturation, gamete release synchrony and duration of the spawning season. Changes in sea surface temperature (Keith et al. 2016), solar and moonlight irradiance (Jokiel, Ito & Liu 1985, Penland, Kloulechad, Idip, & Van Woesik 2004), periods of darkness (Lin et al. 2021, de la Torre Cerro et al. 2025), tidal cycles (Gouezo, Doropoulos & Fabricius 2020), or regional wind patterns (Sakai et al. 2020, van Woesik 2010) have been related to spawning processes. Yet, these interactions are not fully described. Recent findings also suggest that global change and anthropic activities may modify previous patterns in spawning synchrony, hence compromising coral reproduction (Shlesinger & Loya 2019, Davies et al. 2023). However, consistent long-term and large-scale data are still missing to accurately quantify these changes. Despite all observations available to date (e.g. Coral Spawning Database, the first global spawning database initiative: Baird et al. 2021), predicting spawning seasons and single events for most coral species remains difficult (Baird et al2022, Komoto, Lin, Nozawa & Satake 2023). Furthermore, the scarcity of data has given birth to a tangle of conflicting theories on the role of lunar cycle or temperature effects on spawning synchrony (Keith et al. 2016, Komoto, Lin, Nozawa & Satake 2023, van Woesik, Lacharmoise & Köksal 2006).

The majority of coral reproduction studies focus on night-time coral spawning (Baird et al. 2022) while daytime spawning corals have received far less attention (Gouezo, Doropoulos & Fabricius 2020). Because night-time spawning is challenging to study due to observers’ schedule and logistics constraints, scientists have used proxies to assess spawning time, such as the collection of coral samples to examine gamete maturation under the microscope, and laboratory transplant experiments with artificial light and temperature regimes (Henley et al. 2022, Howells, Abrego, Vaughan & Burt 2014). However, these invasive substitutes to *in situ* observations can lead to discrepancies in spawning times compared to natural conditions, and are technically challenging and costly. *In situ* consistent long-term, real-time and species-specific observations at a variety of geographical scales to document spatially synchronised spawning are nearly non-existent.

Taxon-wise, *Acropora* (Oken, 1815) is by far the most intensively coral genus studied for spawning (Baird et al. 2022). Studies on other coral taxa are emerging but remain scarce and are even more data-limited (Gouezo, Doropoulos & Fabricius 2020, Baird et al. 2021). This includes *Porites rus* (Forskål, 1775), for which infrequent and brief descriptive studies have been available only recently (Bronstein & Loya 2011, Gouezo, Doropoulos & Fabricius 2020). *Porites rus* is a gonochoric spawning coral species (Penland, Kloulechad, Idip & Van Woesik 2004) found in the Indian and Pacific Oceans and in the Red Sea (Terraneo et al. 2021) from the near-surface to more than 65 m deep (Englebert et al. 2017). It colonises all habitats, including fringing reefs, channels and oceanic outer slopes, both in low and high current environments. It is abundant in many reefs of the Indian and Pacific Oceans, commonly forming massive coral heads in lagoons or large plates at greater depths, while branching and encrusting morphs also exist. This morphologic plasticity demonstrates its adaptation to different depths, current strength and light intensity (Jaubert 1977, Forsman, Barshis, Hunter & Toonen 2009, Padilla-Gamiño, Hanson, Stat & Gates 2012). Finally, it is highly resistant to thermal (Lenz & Edmunds 2017), pH (Lenz & Edmunds 2017), sedimentation (Padilla-Gamiño, Hanson, Stat & Gates 2012, Golbuu, Fabricius, Victor & Richmond 2008) and light (Hoadley et al. 2021) stresses. Corals presenting these properties have been commonly considered as ‘winners’ in front of global change, with the potential to supplant other coral taxa in the future (Loya, Sakai, Yamazato & Nakano 2001, van Woesik, Sakai, Ganase & Loya 2011, McWilliam, Pratchett, Hoogenboom & Hughes 2020). Nevertheless, this potential has not been comforted for the majority of these ‘winning’ species by a better knowledge on their spawning spatio-temporal patterns and processes.

To fill this gap, share the findings, foster public interest and organise coral spawning monitoring with citizens, a non-profit organisation associated with a monitoring network was created in November 2021 to systematise a long-term *P. rus* spawning data collection. Citizen science, using appropriate and simple protocols, is an inexpensive yet efficient means of collecting loads of robust observations to fuel scientific datasets (de Sherbinin et al. 2021). With limited financial resources and the participation of more than 300 citizen scientists of all ages, the organisation gathered and assembled, between 2014 and 2023, spawning occurrence data at reefs across more than 15,000 km of the Southern Hemisphere between 0 and 30 m depth (Fig. 1a).

**Fig. 1:**
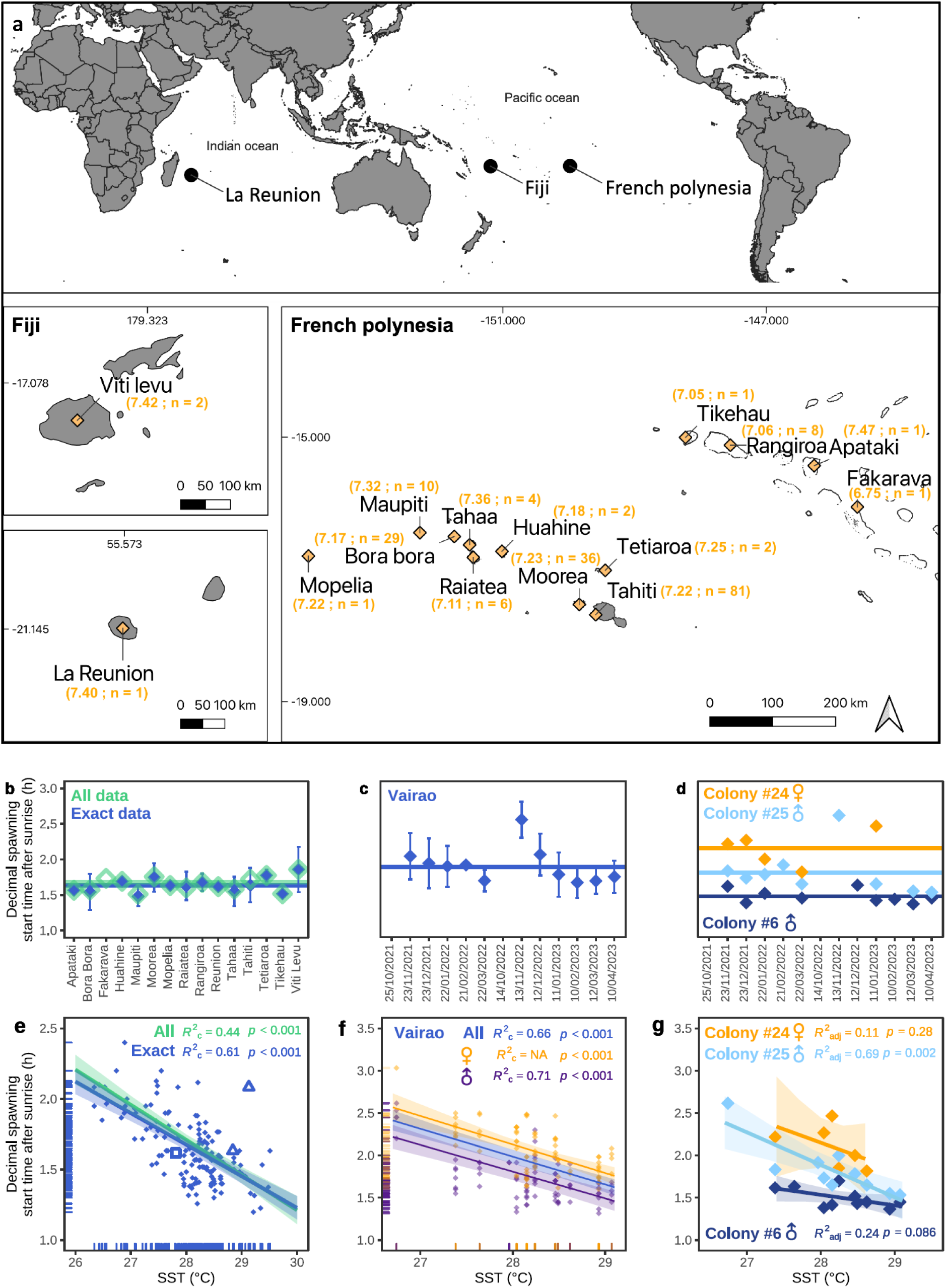
*Porites rus* spawning is synchronous and modulated by temperature. (a) *P. rus* decimal spawning start time (local time in decimal hours) at each island, averaged across shallow sites and spawning occurrences (n) 5 DAFM between 2014 and 2023, based on the time zone corresponding to each location. (b) Mean decimal spawning start time after sunrise (in decimal hours) for all monitored islands between 2014 and 2023 (“La Reunion”: La Reunion Island in the Indian Ocean; “Viti Levu” in Fiji; others: French Polynesian islands) at shallow sites using exact data (blue symbols; SD represented as vertical bars) and all data (exact and approximate data: green symbols; SD not shown for clarity). The horizontal lines represent the decimal spawning start time after sunrise averaged across all corresponding observations (exact and all data). (c) Mean and SD of exact decimal spawning start time after sunrise for shallow colonies (n=30) monitored at Vairao, Tahiti. The horizontal line represents the decimal spawning start time after sunrise averaged across all observations. Note that no spawning occurred in October 2021 and 2022. (d) Exact decimal spawning start time after sunrise for one female and two male colonies monitored in Vairao, Tahiti. No spawning occurred where symbols are absent. The horizontal lines represent the decimal spawning start time after sunrise averaged across the colony observations; note that these two example colonies spawned on averaged earlier than the Vairao average (c). Linear mixed-effect models (LMM) of the relationship between sea surface temperature (SST in °C) and spawning start time after sunrise at large (e: blue symbols and fit: exact data; green fit: exact and approximated data; triangles: Fiji (n=2); square: La Reunion (n=1) data), site (f: orange symbols and fit: females; dark blue symbols and fit: males; blue fit: all data) and colony (g) scales. LMM were fitted to the data with sites (e) and colonies (f) as a random factor. A linear model was fitted to the data in (g). Shading represents the LMM and LM 95% confidence intervals.

This citizen science approach allowed building the largest database ever compiled for *P. rus* spawning. Its analysis provides evidences for recurrent (yearly), highly-predictable (months of the year, day and up to the hour and minute) and large-scale (over 15,000 km) synchronised spawning during daytime for this widely distributed species. The potential cues that influence *P. rus* spawning time are identified, allowing to simply yet accurately predict the months and exact spawning time according to the geographic and environmental characteristics of a site.

## Methods

### Reef scale spawning observations

The organisation developed an *in-situ*, non-invasive observation protocol allowing anyone to observe *Porites rus* spawning at any location in the Southern Hemisphere where *P. rus* is present (Fig. 1a). Observation sites comprising *P. rus* colonies should be around 50 x 50 m large to ensure that all colonies are visible and can be reached rapidly when swimming from one end to the other. Site GPS coordinates or exact location on a map must be recorded. At a specific day of the month and time after sunrise calculated by the organisation, the observer enters into the water 10-15 minutes before the estimated spawning time, wearing a waterproof watch set up to the exact local time and recording the observation date (yyyy-mm-dd). The exact time (hh-mm) at which the first colony on the site is observed to spawn is recorded. If spawning has already started on the site when the observer enters the water, or if the observer is not confident in his watch or observation accuracy, the spawning time recorded is categorised as “approximate”. The time when the last colony of the site finishes to spawn is recorded, and categorised as “exact” or “approximate”. In case no colony is observed to spawn on the whole site, the absence of spawning is recorded.

Following this protocol, data were gathered between 2014 and 2023 by >300 observers and transferred to the organisation by e-mail or directly on a dedicated mobile application. The organisation proceeded to a data check-up before validating each observation for integration into a reef-scale (or site) dataset compiled for the purpose of this study. Initial advice on data gathering were kindly provided by the Coral Spawning Database managers (Baird et al. 2021). Observations from 2014 to 2019 were performed between 0 and 5 m depth, and, because *P. rus* also colonises greater depths (Englebert et al. 2017), from 2020 the monitoring was extended to deeper reefs (6-30 m) using SCUBA.

### Individual colony spawning observation

To better understand spawning time variability at the individual level, 115 *P. rus* colonies were tagged and monthly monitored from 2021 to 2023 (two spawning seasons: 2021-2022 and 2022-2023) in selected sites at 1-2 m depth: 30 colonies in Vairao, 25 in Venus Point and 30 in Arue (Tahiti Island), 10 in Maatea (Moorea Island), and 20 in the Educative Marine Area (Bora Bora Island). Colony monitoring was similar to site monitoring described above, but spawning start and end times were recorded for each tagged colony separately. Since gonochoric coral colonies may sometimes exhibit hermaphroditic behaviour (Neely et al. 2018), sex was checked individually by experienced observers at each spawning event. These data were gathered in a colony-scale dataset.

### Photosynthetic active radiation (PAR) recording

Light effect has been evidenced for other coral species (daytime or night-time spawners) and complex theories involving diurnal light cycles, period of darkness after sunset, lunar photoperiod, nocturnal illumination or twilight colours inducing gamete release were proposed, without any consensus reached to date (Mendes & Woodley 2002, Babcock et al. 1986, Sweeney, Boch, Johnsen & Morse 2011, Boch et al. 2011, de la Torre Cerro et al. 2025). Because *P. rus* lives at different depths characterised by different light regimes, we investigated whether *P. rus* needed to receive a certain quantity of light to induce gamete release (Vize, Hilton, Brady & Davies 2008, Brady, Hilton & Vize 2009, Bouwmeester et al. 2023), as shown with moonlight for night-time spawning species (Jokiel, Ito & Liu 1985, Boch et al 2011, Rosenberg et al. 2017).

To test this, two Odyssey® photosynthetic irradiance (Photosynthetic Active Radiation (PAR): 400-700 nm) loggers were installed on the 12^th^ November and 26^th^ November 2022 at, respectively, 2 and 21 m depth near *P. rus* colonies at the Vairao monitoring site, in an unshaded area. They were calibrated to record PAR irradiance at ten-minute intervals during the spawning season. Loggers were removed on the 4th March and 10^th^ April 2023, respectively. For each spawning date, we cumulated the irradiance values monitored every 10 minutes between sunrise (i.e. just after dark when irradiance becomes >0) and spawning time for the shallow and deep colonies, then averaged the spawning start time and the cumulated irradiance per date.

### Sea surface temperature data and monitoring

Because the spawning season takes place during the Southern Hemisphere warm season (tropical spring and summer), we hypothesised that an increase in sea water temperature along the season may modulate spawning time. Previously, the literature reported the increase in sea surface temperature (SST) as a trigger for the onset of the spawning season (Keith et al. 2016, Gouezo, Doropoulos & Fabricius 2020, Osman et al. 2024). Here, we investigated the impact of temperature on spawning time at a daily scale (i.e. at the exact days of spawning) throughout the season as seawater temperature warmed up, to better understand spawning time variability across months and improve the monthly spawning time predictions.

Daily SST at high resolution (0.05° x 0.05°) for all sites and dates recorded in the reef-scale and colony-scale datasets was collected from the OSTIA (Good et al. 2020) global sea surface temperature reprocessed product database on Marine Copernicus (https://marine.copernicus.eu). To validate the use of these oceanic SST data for lagoon sites, lagoon *in-situ* temperature data were collected from loggers (multiparameter probe NKE model Smatch and Sambat and temperature probe Seabird SBE 56, installed successively at the same location at 5 m depth) located near the Vairao monitoring site: OSTIA SST and *in-situ* temperature data showed a significant correlation (*RMSE*=0.21, *R^2^*=0.95, *p*<0.001: see Appendix S4 Fig. S4.1 in Supporting Information), showing that oceanic OSTIA SST were reliable values to use in our analysis. OSTIA SST were therefore subsequently assigned to each spawning date and location in the reef-scale and colony-scale spawning datasets to assess the effect of spawning day SST on start time after sunrise throughout the spawning season, at different spatial scales and for both colony sexes.

### Statistical data analysis

We fit a linear model (LM) to the exact spawning time for each habitat (shallow lagoon and deep channel-outer slope habitats) as a function of time of sunrise to show the consistency of spawning time relative to the time of sunrise and the lag between spawning start time in the two habitats. We then fitted two LM (one for “exact” only data, and one for “all” data, i.e. approximate and exact data) between depth and spawning start time after sunrise to estimate how the spawning start time after sunrise changes with depth. The adjusted *R*-squared and the *p*-values were calculated.

Linear mixed-effects models (LMM) with site or colony as a random effect were used to study the relationships between SST (fixed effect) and spawning start time after sunrise at different spatial scales (Southern Hemisphere and reef scale), while LM (no random effect) were used to study this relationship at the colony scale. Corresponding conditional and adjusted *R*-squared and *p*-values were calculated.

All analyses were run using R 4.0.3 (R Core Team 2020) and the packages ‘raster’ (Hijmans et al. 2022), ‘lme4’ (Bates et al. 2015) and ‘lmerTest’ (Kuznetsova et al. 2017). Maps were constructed on QGIS 3.16.

## Results

### Citizen science reveals *Porites rus* spawning schedule and large-scale spatial synchrony

*Porites rus* appeared to follow a highly synchronised spawning pattern across different scales (Fig. 1). Opportunistic observations of *P. rus* spawning on 11^th^ November 2014, 8^th^ November 2017 and 27^th^ November 2018 in Moorea and Tahiti (French Polynesia) and the only available publication (Bronstein & Loya 2011) allowed to identify the possible spawning day and time. Accordingly, on 17^th^ November 2019, four sites on two French Polynesia islands were synchronously surveyed using the citizen science approach described in the Methods, collecting data which confirmed the predicted and reported (Bronstein & Loya 2011) spawning time accuracy and synchrony: 5 days after full moon (DAFM) ca. 1.5 to 2 hours after sunrise in fringing reefs (i.e. 1-5 m depth) (Fig. 1a). Total lagoon observations collected between 2014 and 2023 revealed that spawning day and start time (local time and start time relative to sunrise) were extremely synchronised over 15,000 km (Fig. 1a), at 104 reefs spread on the 15 islands sampled (Fig. 1a,b: from Reunion Island in the Indian Ocean to Fakarava Atoll in the Pacific Ocean: mean_exact_=1.64, min_exact_=1.20, max_exact_=2.40, SD_exact_=0.23 decimal hours after sunrise), as well as for all colonies tagged at the monitoring sites (e.g. Vairao, Tahiti: Fig. 1c: mean=1.89, min=1.32, max=3.03, SD=0.33 decimal hours after sunrise; other sites: see Appendix S1 Fig. S1.1a-d).

To explore the duration of the phenomenon within a year, sites and colonies monitored across 9 months (exception: July, August and beginning of September) from 2020 to 2023 evidenced that *P. rus* spawning season extends from October at the earliest to April at the latest, at the same time of the day each month (Fig. 1c,d and Appendix S1 Fig. S1.1). No spawning was detected in May, June and late September at any of the monitored sites, and the season was shorter locally (e.g. no spawning in October in Vairao, Tahiti Island: Fig. 1c). Individual colonies spawning time was highly consistent across months and seasons (Fig. 1c and e.g. three tagged colonies in Vairao, Tahiti: Fig. 1d; other sites: see Appendix S1 Fig. S1.1e-h). Interestingly, we did not detect any change in mean spawning time between the monitored seasons (one-way ANOVA on mean monthly exact spawning time after sunrise for factor “season” with levels “2020-2021”, “2021-2022” and “2022-2023”: *F*(2)= 2.094, *p*=0.154). Data analyses of the two datasets therefore showed a strong consistency in spawning time after sunrise 1) at transoceanic, reef and colony spatial scales, 2) during the whole spawning season and 3) across seasons (Fig. 1b-d).

Monitoring specific colonies in Tahiti, Moorea and Bora Bora allowed characterising gamete release duration precisely, a phenomenon lasting 30.6 minutes on average (SD±13.2 minutes, males and females combined). Additionally, although we focused our observations on the 5^th^ DAFM, we observed that several colonies spawned on the 6^th^ DAFM during the 2022-2023 season at the same scheduled time. Some of these colonies were identified to spawn both on the 5^th^ and on the 6^th^ DAFM, while others spawned only on the 6^th^ DAFM.

Using citizen science data may imply some bias in data accuracy (Jamodiong et al. 2018). Despite a variability in the precision of observed spawning time, the use of both approximate and exact data in the analyses did not result in significant differences in mean spawning time between the “exact only” and “all” data groups (Student’s t-test: *t*(332.74)=-1.5884, *p*=0.1131; Fig.1b).

### Depth and light effects on spawning time

When assessing *P. rus* spawning time at greater depths, unsuccessful attempts through trial and error in Tahiti, Moorea and Takapoto Atoll (French Polynesia) between 7 and 9 a.m. were made, until a coincidental observation on 23^rd^ November 2021 on Tikehau Atoll (French Polynesia) forereef at 15 m depth confirmed spawning at ca. 10 a.m local time. Subsequent attempts to observe spawning at deep sites were successfully scheduled around 10 a.m. in deep lagoons, passes and forereefs.

Colonies in deeper habitats (lagoon channels and forereefs) spawned on average 3.25 hours after shallow colonies (Fig. 2a). Deviations from the linear model did exist, but we did not detect outliers in any of the habitats (Fig. 2a). Similar linear relationships with depth were found when using either the “exact” or the “all” data groups (*R^2^*=0.88, *p*<0.001 and *R^2^*=0.82, *p*<0.001, respectively: Fig. 2b). These results indicate that *P. rus* was extremely well synchronised with sunrise depending on its depth. These highly significant linear relationships can be used to predict the spawning start time after sunrise accurately for sites located at different depths (see Appendix S5), although more work is required to better characterise the spawning time variability observed around 15 m depth (Fig. 2b).

**Fig. 2:**
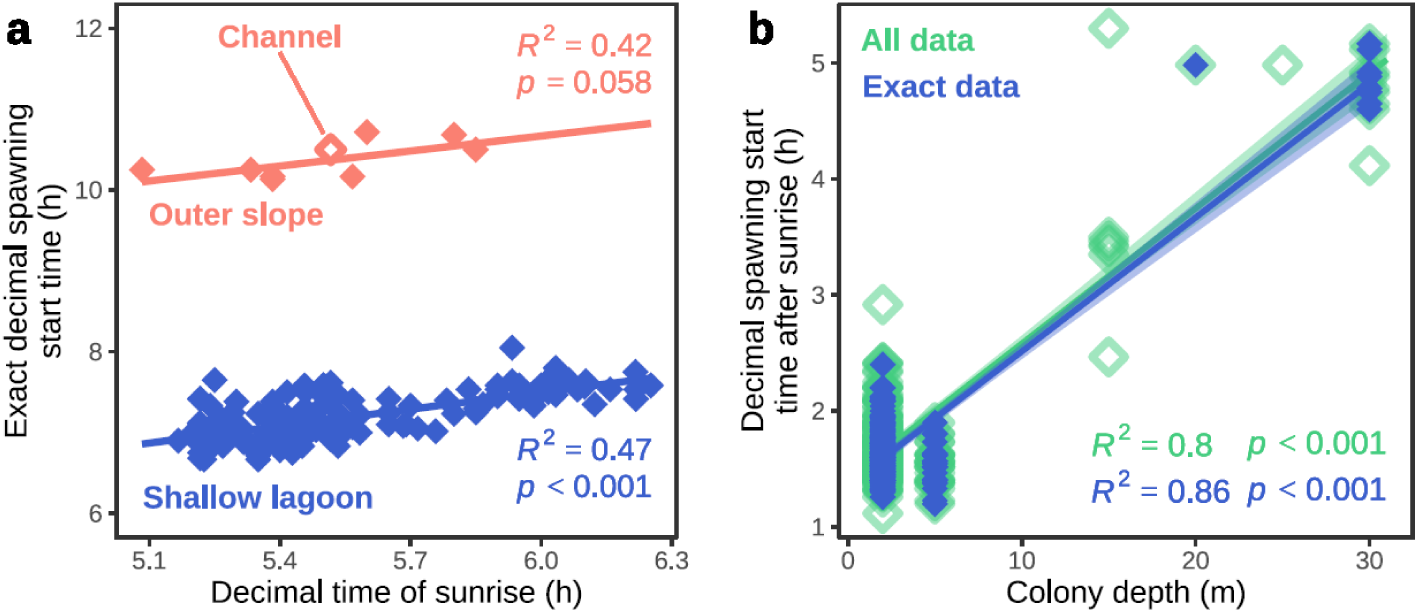
Spawning time varies with depth. Linear models of the relationship between (a) decimal time (in hours) of sunrise and exact decimal spawning time per habitat (“Outer slope” and “Channel” data, located at similar depths, are combined in the same model) and (b) reef location 5 metre-depth bins and spawning start time after sunrise using exact (blue symbols) and all (exact and approximate: green symbols) data. Shading represents the LM 95% confidence intervals.

Since the intensity of light decreases with water depth, we hypothesised that the delay in the onset of spawning at greater depths (Fig. 2a,b) resulted from the diminished intensity of penetrating light, causing the gamete release to be postponed until the colonies had accumulated an adequate amount of light. Accordingly, we observed that: 1) mean spawning time was on average earlier when cumulated irradiance was high (e.g. January and February 2023); 2) the first spawning time of the season (November 2022) occurred later than the others, requiring higher irradiance (a delay that is systematically observed every year at each site, e.g. Fig 1c); and 3) spawning at the deep site started only when the colonies received nearly as much cumulated irradiance at that depth as the shallow colonies, i.e. 5 hours against 1.78 hours after sunrise, with 11017 cumulated W.m^-2^ against 12098 cumulated W.m^-2^, respectively (Fig. 3).

**Fig. 3.**
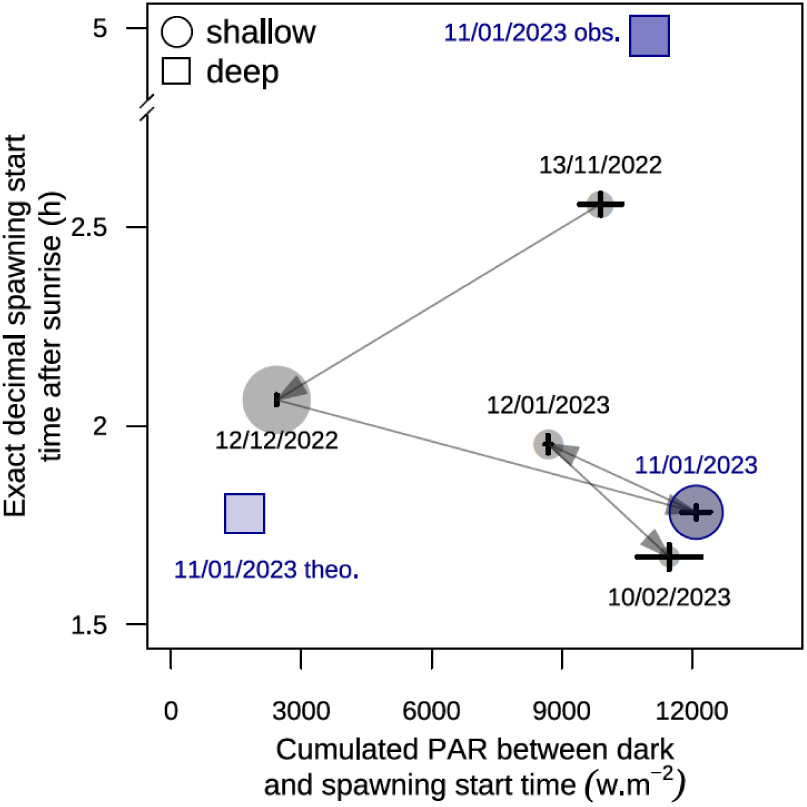
Cumulated photosynthetic active radiation (PAR) irradiance (W.m^-2^) and exact spawning time after sunrise (in hours) for *Porites rus* colonies in Vairao (Tahiti Island) during the 2022-2023 spawning season at low and high depths. Arrows indicate the temporal trajectory for shallow colony spawning, and the spawning date is indicated near each symbol; black bars in each symbol indicate horizontal and vertical SE. Spawning on 11/01/2023 for shallow and deep colonies is highlighted in blue; “obs”: spawning observed for the deep colony; “theo”: theoretical cumulated PAR irradiance if the deep colony had to spawn at the same time as the shallow colonies on this day.

Finally, we calculated the cumulated irradiance for the deep colonies if they had to spawn at the same time as shallow colonies, i.e. 1.78 hours after sunrise. This theoretical irradiance was under the mean lower limit observed during the season (on day 12/12/2022), and nearly 10 times lower than the irradiance received at that date by shallow colonies (on day 11/01/2023) (Fig. 3).

### Temperature and spawning time relationship across scales

A strong significant linear relationship revealed that spawning start time after sunrise occurred earlier at higher SST across oceans (LMM on exact data: *p*<0.001, *R^2^* =0.61, Fig. 1e), for colonies within a site (LMM on Vairao colonies: *p*<0.001, *R^2^*=0.66, Fig. 1f) and for individual colonies (LM on individual colonies, Fig.1g). This indicates that *P. rus* spawned, on average, earlier after sunrise at warm locations, and that spawning occurred earlier at a given location and for single colonies when the ocean warms up along the spawning season (tropical spring and summer) (Fig. 1c,d).

### Synchrony between males and females

Gonochoric *P. rus* male spawning resembles a white opaque cloud of smoke (Fig. 4c) while female spawning is similar to a transparent cloud of <1mm round particles going toward the surface (Fig. 4d). Sites naturally varied in their proportion of male and female colonies with an average of 68±20% males and 32±20% females per site among the tagged colonies. In Vairao, the proportion of colonies that spawned was well-balanced at most months (Fig. 4a) and females started to spawn 19.8 minutes after males (1.72±0.28 hours after sunrise for males, 2.05±0.28 hours after sunrise for females; Student’s t-test *t*(148.67)=-7.11, *p*<0.001, Fig. 4b; other sites: see Appendix S3 Fig. S3.1). Data distribution also indicated that female spawning started during the male spawning peak, then spawning reduced rapidly for males and progressively for females (Fig. 4b). Additionally, both male and female spawning start times were strongly and similarly correlated with SST, from the reef scale (Fig. 1f) down to the individual scale (Fig 1g).

**Fig. 4.**
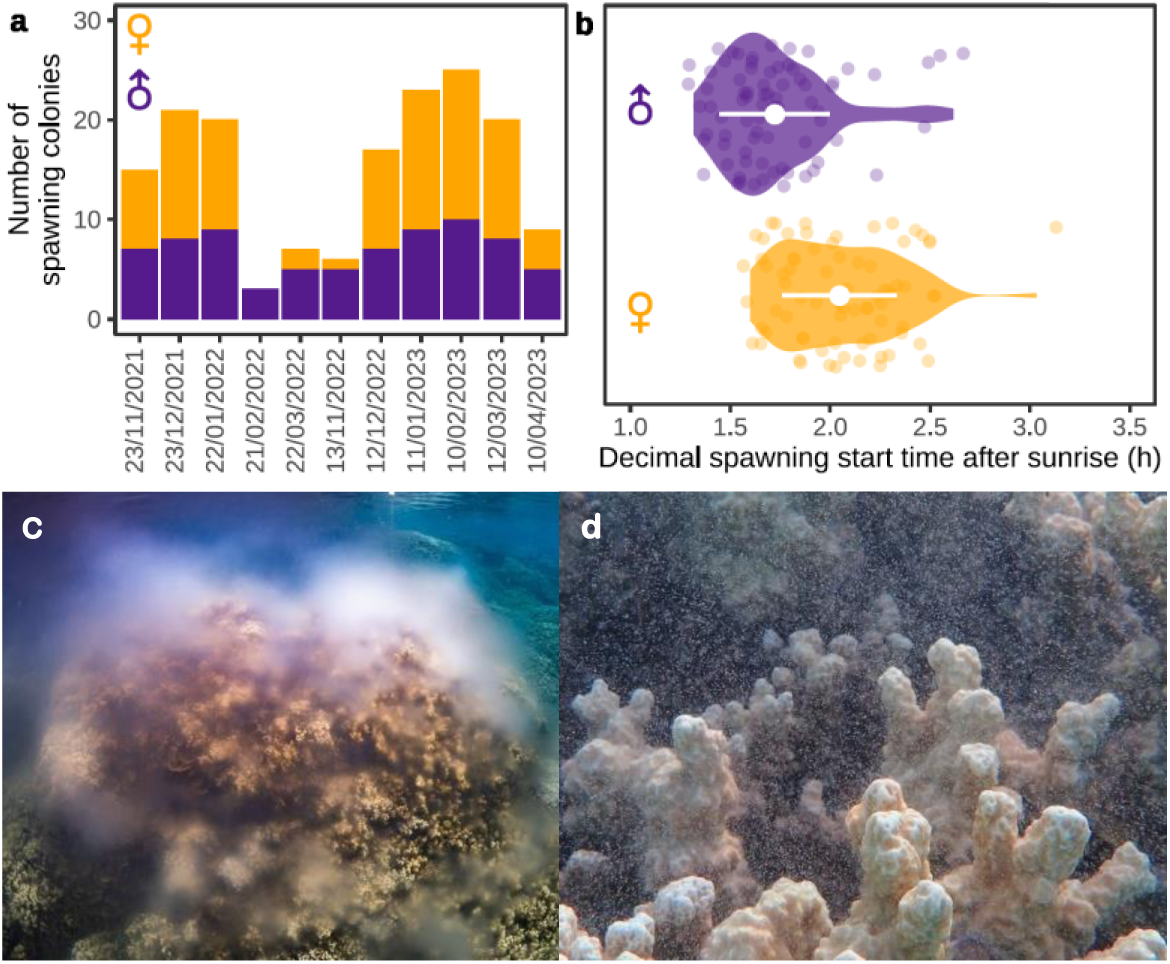
Spawning observations and time difference between *Porites rus* male and female colonies. (a) Number of tagged male and female colonies spawning in Vairao, Tahiti Island, between 2021 and 2023. (b) Decimal spawning start time after sunrise (in hours) at Vairao site for male and female colonies. White dots indicate the mean start time after sunrise and horizontal white lines show SD. Females at Vairao spawn on average 19.8 minutes after males. Male (c) and female (d) spawning.

## Discussion

We showed that *P. rus* spawning occurs monthly from October to April in the Southern Hemisphere at predefined days and times, an unprecedented spawning season duration for a coral (Baird et al. 2022). While coral spawning was known to vary within a range of days (e.g. between the 1st and 11th DAFM: Levitan et al. 2011, Lin & Nozawa 2017) and some coral species were observed to spawn over consecutive months or days at the reef level (Lin & Nozawa 2017, Jamodiong et al. 2018), the literature remains evasive on the capacity of a single colony to spawn over consecutive months and days. Our colony-scale data analysis revealed that *P. rus* single colonies can spawn for up to 5 consecutive months, which has not yet been reported to date for an entire single colony, neither for *P. rus* or for other coral species (Baird et al. 2022, Gouezo, Doropoulos & Fabricius 2020). Additionally, we showed that *P. rus* colonies have the potential (i.e. the gamete storage capacity and the energy to produce and expulse them) to spawn two days in a row. This consecutive spawning should be the focus of further work to assess whether some colonies fail to capture the lunar cycle cue and spawn only after the 5^th^ DAFM, or if all individuals have the capacity to spawn several days in a row. The monthly and daily recurrence of colony spawning within a single season highlights *P. rus* strong reproduction efficiency, contrary to other corals (Baird et al. 2022), and could explain its large-scale distribution and persistence in stressed environments. Finally, we accurately quantified the delay between male and female spawning, which has been evidenced for other gonochoric coral species and is supposed to increase fertilisation success rate (Vize et al. 2005).

Animal reproduction is known to be largely influenced by light and temperature (Beltran-Frutos, Casarini, Santi & Brigante 2022, Dougherty et al. 2024), two factors responsible for gamete maturation and expulsion in corals (Nozawa 2012, Crowder, Liang, Weis & Fan 2014, Wuitchik et al. 2019, Komoto, Lin, Nozawa & Satake 2023). Here, we showed that the increase in temperature during the spawning season (tropical spring and summer) induced earlier spawning at large scale as well as for single colonies, highlighting that temperature influences coral physiology throughout the season, probably resulting from chemical reaction velocity increase in reproductive organs and cells at high temperatures (Crowder, Liang, Weis & Fan 2014) or from a thermally-variable regulation of the expression of genes associated with reproduction (Wuitchik et al. 2019). A relationship with temperature has been evidenced at two sites in Japan (Lin & Nozawa 2023), but the present results also hold for single colonies as much as for sites situated 15,000 km apart. Furthermore, similar temperature-spawning time relationships for males and females suggest that both sexes undergo similar temperature regulation processes throughout the season. Light, on the other end, acts as a trigger for gamete expulsion. However, this has mainly been highlighted for night-time spawning coral species with moonlight (Komoto, Lin, Nozawa & Satake 2023). Despite our PAR logger data was limited to one season, the daylight-spawning time relationship found here is encouraging to extend monitoring of individual colonies at intermediate and high depths to gather additional data to support the cumulated PAR irradiance hypothesis for daytime spawning corals, and analyse the effects of the combination of both temperature and light on spawning time variability.

Interindividual (i.e. among colonies) variation in spawning time and date (e.g. 6^th^ DAFM) does exist at the individual colony scale. This natural intraspecific variability may constitute a means of increasing spawning duration at the reef scale to favour fertilisation, genetic mixing and genetic variability (Levitan et al. 2011, Oomen & Hutchings 2015). More research is however needed to understand this variability and correlate it to the geographic (e.g. depth, latitude and proximity with the equator) and environmental (e.g. light, temperature, sun and moon, turbidity, salinity, tide) factors. The physiological and genetic processes (Celis et al. 2016) inducing gamete maturation and synchronised release should also be investigated to assess whether spawning is induced individually by environmental cues, genetic precision and/or conspecifics molecular signalling within a site. This could provide additional insights into whether certain colonies in specific locations exhibit a regulated delay in spawning or, alternatively, a dysfunction in their spawning physiology due to genetics. Despite these variability sources, the overall spawning phenomenon of *P. rus* occurring over thousands of kilometres is highly synchronous and follows definite cues that we have partially identified.

This study provides evidence that well-organised citizen science can contribute to accumulating valuable and accurate data to study large-scale biological phenomena (de Sherbinin et al. 2021). Similar patterns were found for exact-only and all spawning data, supporting the validity of using unexperienced volunteers to collect reliable scientific observations. Monitoring sites in many Indo-Pacific islands are effortlessly accessible, and the spawning window at the beginning of the day can meet anyone’s agenda (see Appendix S5). A bias regarding the exact time of spawning does exist at intermediate depths (ca. 15 m), but we showed here that it can be largely thwarted by the accessibility of the extremely simple protocol (see Material and Methods and Appendix S5) which attracts a large number of observers, and thus data, at large-scale and at no cost. As an example, the largest synchronous spawning in the database was observed on 23^rd^ December 2021 by 115 volunteers recording 30 simultaneous spawning incidences at 32 reefs over 9 islands spread over 360 km. Gathering the same amount of data with classical scientific means would have been impossible without consistent funding.

Predicting coral spawning is crucial to limit the impacts of coastal development on coral reefs. Understanding the spawning processes and triggers of a wide-spread species can precisely inform decision making and spatial management at local to large scales. Despite other few countries (e.g. Australia: Styan & Rosser 2012) have been following well-established precautionary rules for decades, they still face the lack of accuracy in predicting the exact day of coral spawning. In the case of *P. rus*, which consists in a large proportion of the fringing coral community in many tropical countries, spawning is sufficiently predictable for clear precautionary guidelines to be set up and followed. Locally, the organisation is actively encouraging coastal and marine construction works not to alter the marine environment (e.g. no digging, no mining, no river curation, no turbulences, no underwater noise disturbance, etc.) during the spawning days across the entire, highly predictable spawning season. Alerting about the fragility of coral reefs, especially in the fringing reef area, is crucial and has proven to be efficient, as local construction companies are now following guidelines and asking for environmental impact advice before implementing their works. Raising awareness about environmental fragility by engaging the public through field observations can be an effective and direct way to protect the environment before new, appropriate laws are enacted and enforced by governments.

## Conclusion

In the context of global coral reef degradation, our findings bring significant input to foster coral biological clock research. Global change is suspected to alter synchronic ecological processes at large scale (Hacket-Pain & Bogdziewicz 2021, Karatayev, Munch, Rogers & Reuman 2025), including coral spawning synchrony (Shlesinger & Loya 2019). So far, we have observed no change in *P. rus* spawning time over the years. This constancy in spawning synchrony may be due to *P. rus* high resistance to environmental and anthropic stresses (Golbuu, Fabricius, Victor & Richmond 2008, Padilla-Gamiño, Hanson, Stat & Gates 2012, Lenz & Edmunds 2017, Hoadley et al. 2021), and accumulating data over additional years will help determine whether the synchrony of *P. rus* gamete release remains stable in the future. The absence of detecting long-term changes could mean that *P. rus* is insensitive to environmental changes or able to adapt its reproductive processes to these changes, hence procuring an additional characteristic of a “winner” species in front of global change. We therefore believe that more attention should be given to *P. rus* as it is an accessible coral that has long been overlooked. Despite the study of *P. rus* spawning is only in its early stages here, our results on the spawning time relationship with depth, light and temperature at various spatial and time scales can catalyse research for *P. rus* around the globe in several scientific fields, such as biogeography, macroecology, physiology and genetics, to disentangle the intrinsic and extrinsic cues regulating internal clocks leading to gamete maturation and release synchrony.

## Data availability

The data supporting the findings of this study are provided in the supplementary material of this article.

## Code availability

The code generating the analyses is freely available upon request to C.M. It can be provided to GEB during the evaluation of the manuscript if required.

## Notes

### Competing Interest Statement

The authors have declared no competing interest.

