## Supplementary Material for "Shining a light on daytime coral spawning synchrony across oceans"

### Appendix S1

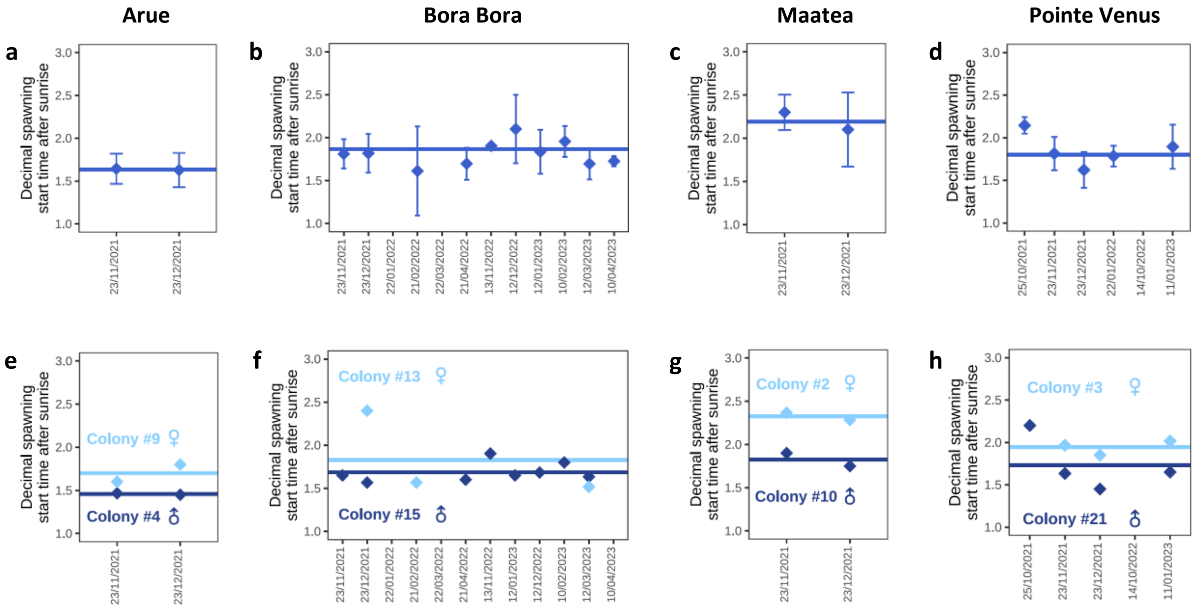


**Fig. S1.1. Reef and colony scale spawning time.** Spawning start time after sunrise (in decimal hours) for colonies tagged at Arue Tahiti (a), Bora Bora (b), Maatea Moorea (c) and Pointe Venus Tahiti (d), and for some single colonies at these same sites (e-h) across seasons. Lines represent the mean, and vertical bars at reef scale, the SD.

**Appendix S2**

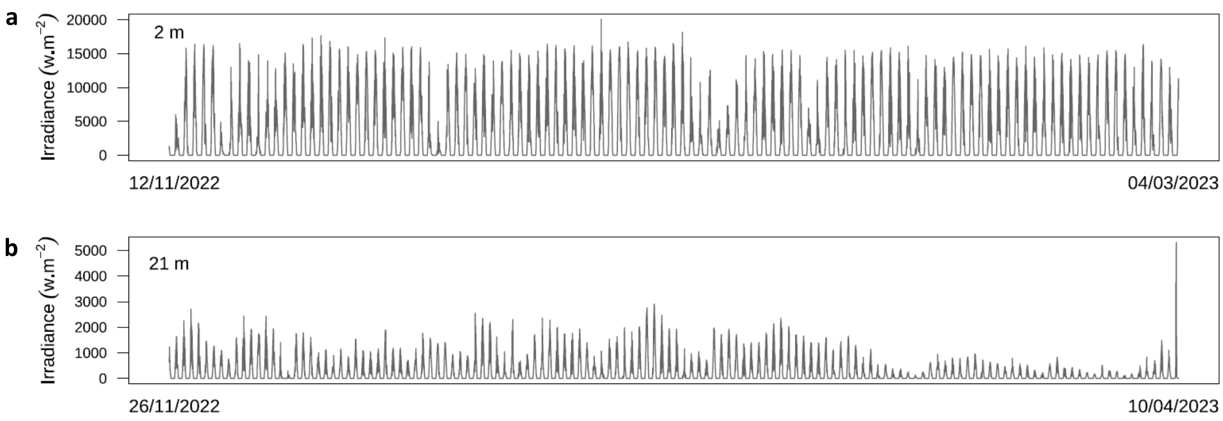


**Fig. S2.1. Irradiance.** Photosynthetic active radiation (400-700 nm) irradiance (W.m$^{-2}$) measured every 10 minutes at the Vairao site at 2 (a) and 21 m depth (b) near monitored *P. rus* colonies during the 2022-2023 spawning season.

**Appendix S3**

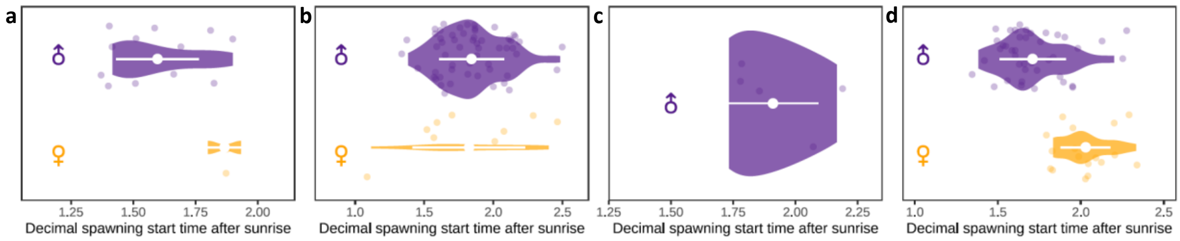


**Fig. S3.1. Male and female spawning.** Spawning start time after sunrise (in decimal hours) for males and females at Arue-Tahiti (a), Bora Bora (b), Maatea-Moorea (c) and Pointe Venus-Tahiti (d).

**Appendix S4**

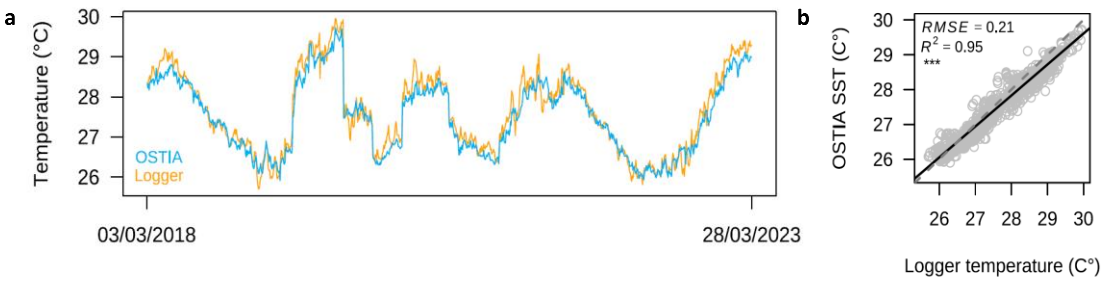


**Fig. S4.1. *In situ* and OSTIA temperature data.** *In situ* temperature from logger (in °C) and sea surface temperature (SST in °C) from OSTIA (Good et al., 2020) global SST reprocessed product covering the data period (a), and correlation between logger temperature and OSTIA SST (b; the dotted line represents the 1:1 line).

### Appendix S5

**Predicting the spawning time to observe *Porites rus* spawning**

To accurately predict *P. rus* spawning time at any Indian and Pacific Oceans reef, six parameters need to be considered and intersected:

**1. Date:** count 5 days after full moon; do not hesitate to look for spawning on the $6^{th}$ day after full moon as consecutive spawning may occur at reef and colony scales.

**2. Time:** check the time of sunrise at your exact location. We suggest using *timeanddate.com*.

**3. Depth:** identify the habitat you are observing and its depth (Fig. 3a, b): this will inform whether spawning will start soon or later after sunrise. Add *ca.* 1h30 after sunrise for colonies located ≤5 m deep and *ca.* 5h after sunrise for colonies located ≥20 m deep. Colonies situated between 5 and 20 m deep should spawn in between.

**5. Month:** As seawater temperature increases along the spawning season, events from December might be advanced a few minutes.

**4. Weather:** Check the weather on the day of spawning: the event is likely to be slightly delayed (a few minutes) if the weather is cloudy because of less light penetrating the water.

**6. Sex:** For single colony observations, identify (using preliminar spawning observations if available) whether your colony is a male or a female: scheduled time given in 3. is for males, and females will start 15-20 minutes later.

**7. Event length:** Spawning of a single colony can last about 30-40 minutes. Spawning at site level (i.e. several colonies) can last longer as some colonies start to spawn later and therefore finish later.

**8. Report** your observations to the organisation mentioning: your name, the day, start and end times of observation in the water, start and end times of spawning, the absence of spawning if no spawning is observed, sex of the colonies if checked individually, location (exact description or GPS coordinates), depth and any additional environmental conditions and comments that you find useful.

Above these recommendations, we have noted that the first spawning event of the season (October or November depending on reefs) can occur later than predicted; do not hesitate to stay longer in the water to wait for the spawning to start. Finally, these predictions can vary according to reefs and colonies due to site specificities and interindividual variability. Therefore, whatever the exact time you predict, we suggest you start the underwater observation 15 minutes before schedule.
